# Sampling strategy optimization to increase statistical power in landscape genomics: a simulation-based approach

**DOI:** 10.1101/603829

**Authors:** Oliver Selmoni, Elia Vajana, Annie Guillaume, Estelle Rochat, Stéphane Joost

## Abstract

An increasing number of studies are using landscape genomics to investigate local adaptation in wild and domestic populations. The implementation of this approach requires the sampling phase to consider the complexity of environmental settings and the burden of logistic constraints. These important aspects are often underestimated in the literature dedicated to sampling strategies.

In this study, we computed simulated genomic datasets to run against actual environmental data in order to trial landscape genomics experiments under distinct sampling strategies. These strategies differed by design approach (to enhance environmental and/or geographic representativeness at study sites), number of sampling locations and sample sizes. We then evaluated how these elements affected statistical performances (power and false discoveries) under two antithetical demographic scenarios.

Our results highlight the importance of selecting an appropriate sample size, which should be modified based on the demographic characteristics of the studied population. For species with limited dispersal, sample sizes above 200 units are generally sufficient to detect most adaptive signals, while in random mating populations this threshold should be increased to 400 units. Furthermore, we describe a design approach that maximizes both environmental and geographical representativeness of sampling sites and show how it systematically outperforms random or regular sampling schemes. Finally, we show that although having more sampling locations (between 40 and 50 sites) increase statistical power and reduce false discovery rate, similar results can be achieved with a moderate number of sites (20 sites). Overall, this study provides valuable guidelines for optimizing sampling strategies for landscape genomics experiments.

## Introduction

Landscape genomics is a subfield of population genomics, with the aim of identifying genetic variation underlying local adaptation in natural and managed populations (Balkenhol et al., 2017; Joost et al., 2007; Rellstab, Gugerli, Eckert, Hancock, & Holderegger, 2015). The approach consists of analyzing genomic diversity and environmental variability simultaneously in order to detect genetic variants associated with a specific landscape composition. Studies of this kind usually incorporate an analysis of population structure, such that neutral genetic variation can be distinguished from adaptive variation (Rellstab et al., 2015). Over the last few years, the landscape genomic approach is becoming more widely used (see Tab. 1; Balkenhol et al., 2017; Rellstab et al., 2015). It is being applied to a range of species, including livestock (Colli et al., 2014; Lv et al., 2014; Pariset, Joost, Marsan, & Valentini, 2009; Stucki et al., 2017; Vajana et al., 2018), wild animals (Harris & Munshi-South, 2017; Manthey & Moyle, 2015; Stronen et al., 2015; Wenzel, Douglas, James, Redpath, & Piertney, 2016), insects (Crossley, Chen, Groves, & Schoville, 2017; Dudaniec, Yong, Lancaster, Svensson, & Hansson, 2018; Theodorou et al., 2018), plants (Abebe, Naz, & Léon, 2015; De Kort et al., 2014; Pluess et al., 2016; Yoder et al., 2014) and aquatic organisms (DiBattista et al., 2017; Hecht, Matala, Hess, & Narum, 2015; Laporte et al., 2016; Riginos, Crandall, Liggins, Bongaerts, & Treml, 2016a; Vincent, Dionne, Kent, Lien, & Bernatchez, 2013).

**Table 1.** Sampling design in landscape genomics studies. A non-exhaustive list of landscape genomics studies, highlighting different species and their related sampling strategies.

\* Numbers from the Vincent et al. report (2013) concerning the *non-pooled* samples.
| Study | Species | Sampling Design (D) | Sampling Locations<br>(L) | Sample Size<br>(S) |
| --- | --- | --- | --- | --- |
| Colli et al. 2014 | Goat | Spatial and breed representativeness | 10 sites | 43 |
| Pariset et al. 2009 | Goat | Spatial and breed representativeness | 16 regions | 497 |
| Stucki et al. 2017, Vajana et al.<br>2018 | Cattle | Spatial representativeness | 51 regions | 813 |
| Harris and Munshi-South, 2017 | White-footed<br>Mouse | Habitat representativeness | 6 sites | 48 |
| Stronen et al., 2015 | Wolf | Opportunistic, population<br>representativeness | 59 sites | 59 |
| Wenzel et al., 2016 | Red Grouse | Spatial representativeness | 21 sites | 231 |
| Crossley et al., 2017 | Potato Beetle | Habitat representativeness | 16 sites | 192 |
| Dudaniec et al., 2018 | Damselfly | Environmental and spatial<br>representativeness | 25 sites | 426 |
| Theodorou et al., 2018 | Red-tailed<br>bumblebee | Habitat representativeness | 18 sites | 198 |
| Abebe et al., 2015 | Barley | Spatial representativeness | 10 regions | 260 |
| De Kort et al., 2014 | Black alder | Spatial and habitat representativeness | 24 populations | 356 |
| Pluess et al., 2016 | European beech | Spatial and environmental<br>representativeness | 79 populations | 234 |
| Yoder et al., 2014 | Barrelclover | Spatial representativeness | 202 sites | 202 |
| DiBattista et al., 2017 | Stripey Snapper | Spatial representativeness | 51 sites | 1,016 |
| Hecht et al., 2015 | Chinook salmon | Spatial representativeness | 53 sites | 1,956 |
| Laporte et al., 2016 | European Eel | Spatial and environmental<br>representativeness | 8 sites | 179 |
| Vincent et al., 2013 | Atlantic Salmon | Spatial representativeness | 26* rivers | 641* |

Sampling strategy plays a pivotal role in experimental research, and must be theoretically tailored to the aim(s) of a study (Rellstab et al., 2015; Riginos et al., 2016). In the context of landscape genomics, the sampling design should cover a spatial scale representative of both the demographic processes and the environmental variability experienced by the study population (Balkenhol et al., 2017; Leempoel et al., 2017; Manel et al., 2010; Rellstab et al., 2015). This is imperative to be able to properly account for the confounding effect of population structure, to provide a biologically meaningful contrast between the environmental variables of interest and to definitely allow the search for actual adaptive variants (Balkenhol et al., 2017; Manel et al., 2010; Rellstab et al., 2015). Consequently, extensive field sampling is generally required and needs to be coupled with high-throughput genome sequencing to characterize samples at a high number of loci (Balkenhol et al., 2017; Rellstab et al., 2015). Beyond these theoretical aspects, pragmatic choices need to be made with regards to financial and logistic constraints that are often imposed (Manel et al., 2010; Rellstab et al., 2015). A sampling strategy is constituted of: i) sampling design (the spatial arrangement of the sampling locations, D); ii) the number of sampling locations (L); and iii) sample size (the number of individuals sampled, N; Tab. 1). The care with which these parameters are defined affects the scientific output of an experiment as well as its costs (Manel et al., 2010; Rellstab et al., 2015).

The landscape genomics community has traditionally focused on formulating theoretical guidelines for collecting individuals throughout the study area. In this literature, particular emphasis has been placed on how spatial scales and environmental variation should be accounted for when selecting sampling sites (Leempoel et al., 2017; Manel et al., 2010; Manel et al., 2012; Rellstab et al., 2015; Riginos et al., 2016). Theoretical simulations have shown that performing transects along environmental gradients or sampling pairs from contrasting sites which are spatially close reduced false discovery rates caused by demographic processes confounding effects (De Mita et al., 2013; Lotterhos & Whitlock, 2015). However, in these studies the environment was described using a single variable, which oversimplifies the choice of sampling sites. In fact, in a real landscape genomics application, several variables are usually analyzed in order to explore a variety of possible environmental pressures causing selection (Balkenhol et al., 2017). The concurrent use of several environmental descriptors also allows to control for the bias associated with collinear conditions (Rellstab et al., 2015). Furthermore, these studies focused on the comparison of different statistical methods with the drawback of confronting only a few combinations of the elements determining the sampling strategy (De Mita et al., 2013; Lotterhos & Whitlock, 2015). Last but not least, the number of samples used in the simulations (between 540 and 1800; Lotterhos & Whitlock, 2015) appear to be unrealistic for use in most of real landscape genomic experiments (Tab.1) and thus the guidelines proposed are scarcely applicable in practice.

For these reasons, there is a need to identify pragmatic and realistic guidelines such that a sampling strategy is designed to maximize statistical power, minimize false discoveries, and optimize efforts and money expenses (Balkenhol et al., 2017; Rellstab et al., 2015). In particular, the fundamental questions that need to be addressed are: i) how to determine the spatial arrangement of sampling locations; ii) how to organize sampling effort (for instance preferring many samples at few sites, or rather fewer samples at many sites); and iii) how many samples are required to obtain sufficient statistical power (Rellstab et al., 2015; Riginos et al., 2016).

In this paper, we investigate how the outcome of landscape genomic analyses is driven by the sampling strategy. We ran simulations using a fictive genetic dataset encompassing adaptive genotypes shaped by real environmental variables. The simulations accounted for antithetic demographic scenarios encompassing strong or weak population structure. We proposed sampling strategies that differed according to three elements: sampling design approach (D), number of sampling locations (L) and sample size (number of samples, N). For each of these three elements, we measured their relative impacts on the analyses’ true positive rates (TPR) and false discovery rates (FDR), as well as their impact on the predictive positive value (PPV; Marshall, 1989) of the strongest adaptive signals.

## Material & Methods

The iterative approach we designed to test the different sampling strategies required that a new genetic dataset encompassing neutral and adaptive variation was created at every run of the simulations. A simulated genomic dataset can be constructed by means of software performing coalescent (backward-in-time) or forward-in-time simulations (Carvajal-Rodríguez, 2008). However, methods using coalescent simulations (for ex. SPLATCHE2; Ray, Currat, Foll, & Excoffier, 2010) did not match our needs as they cannot compute complex selective scenarios (for instance those involving multiple environmental variables; Carvajal-Rodríguez, 2008). We could not use forward-in-time methods either, as they are slow and therefore not compatible with the computational requirements of our simulative approach (Carvajal-Rodríguez, 2008). For these reasons, we developed a customized framework in the R environment (version 3.3.1; R Core Team, 2016) to compute both neutral and adaptive genetic variation based on gradients of population membership and environmental variations, respectively (Fig. 1). Prior to running the simulations across the complete dataset (the multivariate environmental landscape of Europe), we tested our approach on a reduced dataset and compared it to a well-established forward-in-time simulation software (CDPOP, version 1.3; Landguth & Cushman, 2010). This step allowed us to define the optimal parameters required to simulate two types of demographic scenarios: panmictic (no dispersal constraints, random mating) and structured (dispersal and mating limited by distance).

**Figure 1.**
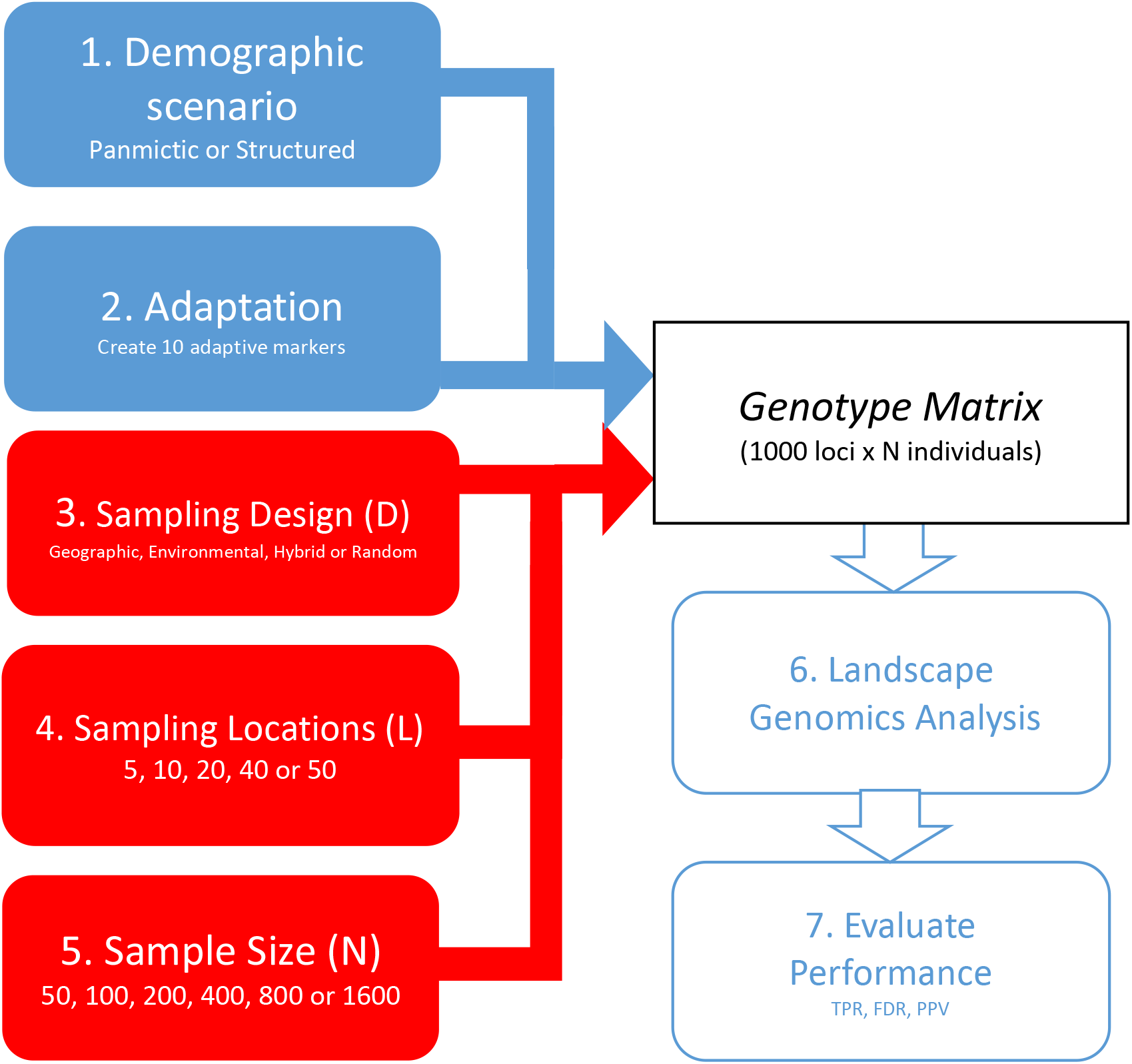
Workflow for each iteration of the simulative approach. The seven steps taken for every iteration. Starting with the blue boxes, the genetic set-up is established by selecting the demographic scenario (panmictic or structured), which determines the neutral structure, and by picking the environmental variables implied in adaptation. The environmental variable of interest and the strength of selection is randomly sampled for each of the 10 adaptive markers. Following this, the sampling strategy (here shown with red boxes) is set as a combination of design approach (geographic, environmental, hybrid or random), number of sampling locations (5, 10, 20, 40 or 50 locations) and sample size (50, 100, 200, 400, 800 or 1600 samples). This results in the creation of a genotype matrix that undergoes a landscape genomics analysis. At the end of iterations, statistical power (TPR) and false discovery rate (FDR) of the analysis and statistical predictive positive value of the strongest associations (PPV) are calculated to assess the performance of the sampling strategy.

We then proceeded with the simulations on the environmental dataset of Europe. At each iteration, a new genetic background encompassing neutral and adaptive variation was computed (Fig. 1 steps 1 and 2). Subsequently, a sampling strategy was applied as a combination of sampling design (D), number of sampling locations (L) and sample size (N) (Fig. 1, steps 3, 4 and 5), resulting in the generation of a genetic dataset that, coupled with environmental data, underwent a landscape genomics analysis (Fig. 1, step 6). At the end of each iteration, three diagnostic parameters were calculated: true positive rate (TPR, *i.e.* statistical power) and false discovery rate (FDR) for the analysis, as well as the predictive positive value (PPV) of the strongest genotype-environment associations (Fig. 1, step 7).

At the end of the simulations, we analyzed how each element of sampling strategy (D, L, N) affected the rates of the three diagnostic parameters (TPR, FDR, PPV) under the two demographic scenarios (with or without dispersal constraints). All scripts and data used to perform this analysis are publicly available on Dryad (doi:10.5061/dryad.m16d23c).

### Environmental data

As a base for our simulations, we quantified the environmental settings of Europe (Fig. S1). We retrieved eight climatic variables from publicly available sources (annual mean temperature, mean diurnal range, temperature seasonality, mean temperature of wettest quarter, annual precipitation, precipitation seasonality, precipitation of warmest quarter and altitude; Tab S1; Hijmans, Cameron, Parra, Jones, & Jarvis, 2005; Ryan et al., 2009). In order to work on a relevant geographical scale (Leempoel et al., 2017) while maintaining an acceptable computational speed, the landscape was discretized into grid cells of 50×50 km, using QGIS toolbox (version 2.18.13; QGIS development team, 2009). This resulted in 8,155 landscape sites. Average values of environmental variables were computed for each cell of the landscape using the QGIS zonal statistics tool.

### Computation of genotypes

For the creation of the genotype matrices, we developed an R-pipeline based on probability functions to compute genotypes from population membership coefficients and environmental values (Box S1). The theoretical fundaments of this method are based on the observation that when the population is structured, neutral alleles tend to show similar spatial patterns of distribution (a feature commonly exploited in Fst outlier tests; Luikart et al., 2003; and principal component analyses of genotype matrices; Novembre et al., 2008). Conversely, when a marker is under selection, its genotypic/allelic frequencies correlate with the environmental variable of interest (this is the basic concept of Landscape Genomics; see Balkenhol et al., 2017). For every iteration, 1,000 loci are computed: 10 are set to “adaptive”, while the remaining 990 to “neutral”. They are computed as follows:

- Neutral markers (Box S1a): a parameter (*m)* is set to define the number of population membership gradients used in the simulations, where higher values of *m* result in more complex population structures. Every population membership gradient is simulated by randomly picking one to five landscape locations to represent the center of the gradient. For each landscape location, the geographical distance to the gradient centers (calculated using the R *dist* function) constitutes the membership coefficient. Next, a linear transformation converts this coefficient (Fig. S1) for each sampling site into the probability of carrying a private allele for the population described (*pA*/*PS*). A second parameter (*c*, Box. S2) define this transformation, with values between 0.5 (random population structure) and 0 (strong population structure). The probability of *pA*/*PS* is then used to draw (using the R-stat *sample* function) the bi-allelic genotype for each individual. This procedure is re-iterated for every neutral locus assigned to a specific population membership coefficient. Each of the 990 neutral loci is then assigned to one of the *m* population membership coefficients (probability of assignment equal to 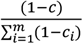 using the R *sample* function.
- Adaptive markers (Box S1b): the probability of carrying an adaptive allele (*pA*/*Env*) is calculated through a linear transformation of a specific environmental gradient. This transformation is defined by two parameters. The first parameter (*s_1_*) determines the amplitude of the transformation, and ranges between 0 (strong selective response) and 0.5 (neutral response; Box S2). The second parameter (*s*_*2*_) shifts the baseline for allele frequencies, and ranges between −0.2 and 0.2 (weakening and strengthening the selective response, respectively; Box S2). Each of the ten adaptive loci are randomly associated with one environmental variable. This implies that some environmental conditions can be associated with several genetic markers, while others with none. For every adaptive locus, the bi-allelic genotype is drawn (using the R-stat *sample* function) out of *pA*/*Env*.

### Evolutionary scenarios and parametrization

Two distinct demographic scenarios were chosen for this study: one involving a population that is not genetically structured (hereafter referred to as the “panmictic population scenario”), and one involving a structured population (hereafter referred to as the “structured population scenario”; see Box S2). In order to define the values of parameters *m, c, s*_*1*_ and *s*_*2*_ that allow the production of these two demographic scenarios, we ran a comparison of our customized simulation framework against simulations obtained using a well-established forward-in-time simulation software for landscape genetics called CDPOP (version 1.3; Landguth & Cushman, 2010).

This comparison was performed on a reduced dataset composed of a 10-by-10 cell grid, covered with two dummy environmental variables extracted from the bioclim collection (Hijmans et al., 2005; Fig. S1a, b). Each cell could host up to 5 individuals, where each individual was characterized at 200 SNPs. In this set-up, we ran CDPOP using two distinct settings: the first that allowed for completely random dispersal and mating movements of individuals (*i.e.* panmictic population scenario), while the second setting restricted movements to neighboring cells using a dispersal-cost based on distance (*i.e.* structured population scenario). In both scenarios, we applied identical mortality constraints related to the two environmental variables, and set for each of them a genetic variant modulating fitness (Fig. S1c, d). Fitness responses were constructed on an antagonistic pleiotropy model (i.e. adaptive tradeoffs, Lowry, 2012), using different intensities to represent moderate (Fig. S1c) and strong selective constraints (Fig. S1d). The following default CDPOP parameters were employed for the remaining settings: five age classes with no sex-specific mortality, reproduction was sexual and with replacement, no genetic mutations, epistatic effects or infections were allowed. The simulations ran for 100 generations and ten replicates per demographic scenario were computed.

In parallel, we ran our customized algorithm to compute genotypes, using the same simplified dataset as above. We iteratively tested all the possible combinations (hereafter referred to as “simulative variants”) of the parameters *m* (values tested: 1, 5, 10, 15, 20, 25), *c* (all possible ranges tested between: 0.1, 0.2, 0.3, 0.4, 0.5), *s*_*1*_ (values tested: 0, 0.1, 0.2, 0.3, 0.4, 0.5) and *s*_*2*_ (values tested: −0.2, −0.1, 0, 0.1, 0.2), and replicated each combination ten times. Following this, we investigated which of the simulative variants provided the closest match with the allele frequencies observed in the CDPOP runs. The comparisons were based on three indicators of neutral structure:

1. Principal component analysis (PCA) of the genotype matrix (Fig. 2a): a PCA of the genotype matrix was performed using the *prcomp* R function for each simulation (of both the CDPOP and the present customized method), where the differential of the variation explained by each principal component was then calculated. When the population is structured, the first principal component usually shows strong differences in the percentage of explained variation compared with the other components (Novembre et al., 2008). In contrast, when the population structure is absent, minor changes in this differential value emerge. The curve describing this differential value was then used for a pairwise comparison between the ten replicates of each CDPOP scenario and the ten replicates of each simulative variant (from the customized method). The curves were compared by calculating the root mean square error (RMSE), then the average RMSE was used to rank simulative variants.
2. F statistic (Fst; Fig. 2b): five areas, which spanned four cells each, were selected to represent subpopulations of the study area. Four areas located at the four corners of the 10-by-10 cell grid and the fifth located at the center. For each simulation, we computed the pairwise Fst (Weir & Cockerham, 1984) between these sub-populations using the *hierfstat* R package (version 0.04; Goudet, 2005). An Fst close to 0 indicates the absence of a genetic structure between sub-populations, while under a structured scenario this value tends to raise (Luikart et al., 2003). The distribution of all the Fst values for the ten CDPOP replicates were compared to the distribution of the Fst of ten replicates of each simulative variant using the Kullback-Leibler Divergence (KLD; Kullback & Leibler, 1951) analysis implemented in the *LaplacesDemon* R package (version 16.1.1; Statisticat & LCC, 2018). KLD was then used to rank simulative variants.
3. Mantel test (Fig. 2c): for each simulation, we computed the genetic and geographic distance between all individuals of the population applying the R *dist* function to the genotype matrix and the coordinates, respectively. Next, we calculated the Mantel correlation (mR; Mantel, 1967) between these two distance matrices using the *mantel.rtest* function implemented in the *ade4* R package (version 1.7, Dray & Dufour, 2007). When mR is close to 0, it indicates the absence of correlation between the genetic and geographical distances, suggesting the absence of genetic structure (*i.e.* panmictic population scenario). In contrast, an mR closer to −1 or +1 indicates that genetic distances match geographic distances, as we would expect in a structured population scenario (Mantel, 1967). The average mR was calculated for each simulative variant and compared to the average mR measured in the two CDPOP scenarios. The resulting difference in mR (ΔmR) was used to rank simulative variants.

**Figure 2.**
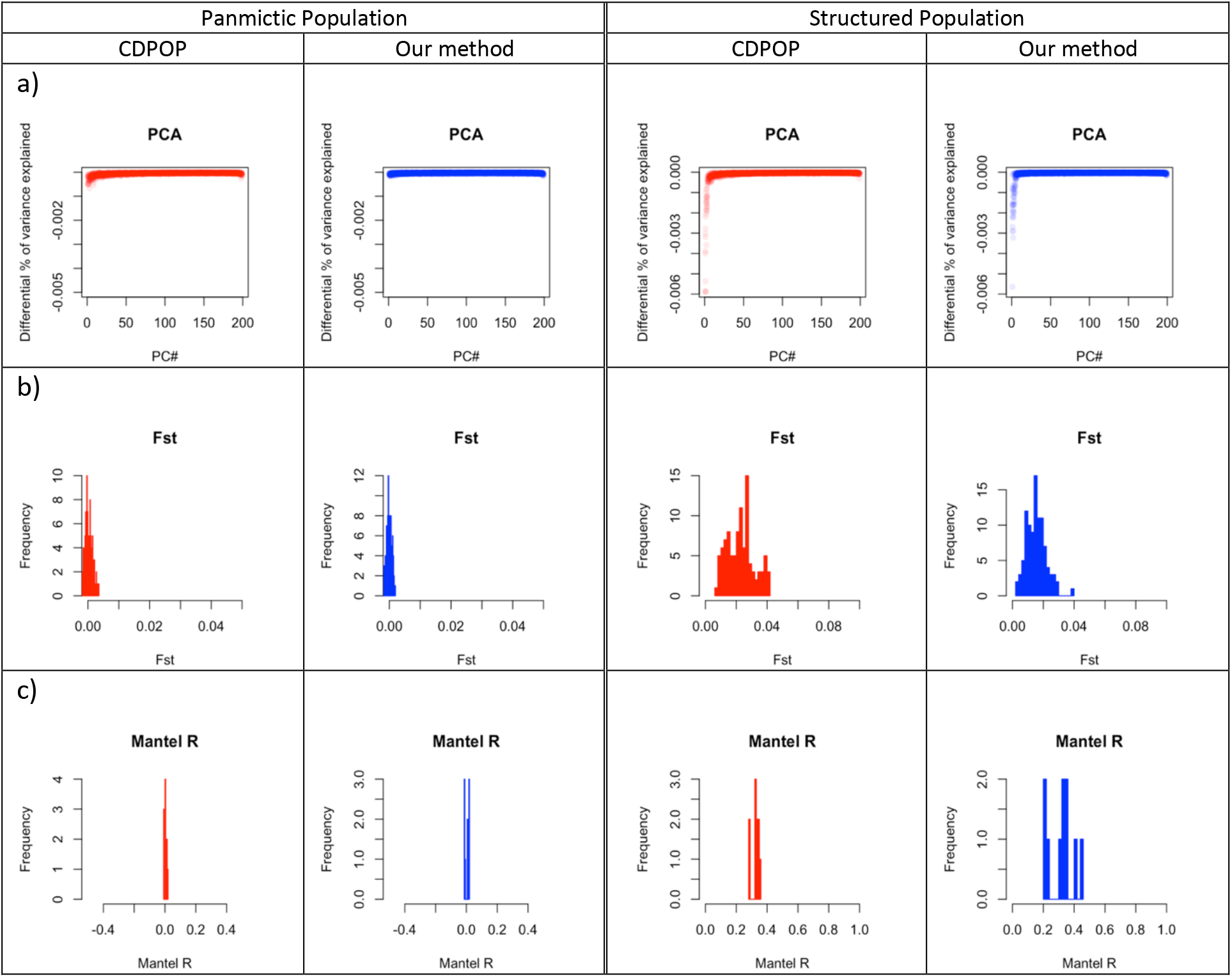
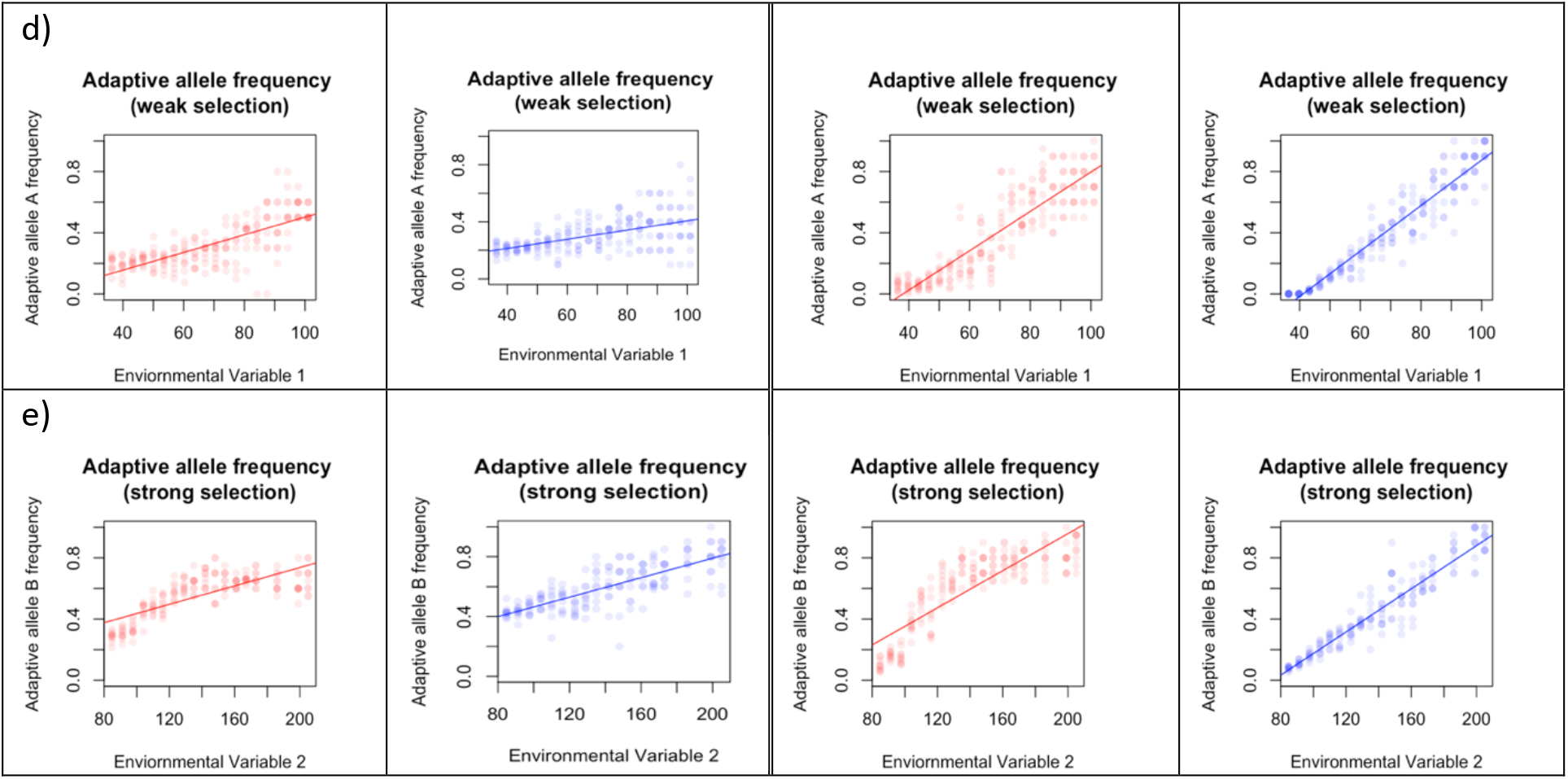
Comparison of genotypes simulated with CDPOP and our method. Two distinct demographic scenarios were conceived, one with random mating (panmictic population) and one with dispersal costs related to distance (structured population). For each of them, CDPOP simulated the evolution of the population over 100 generations (red graphs) and replicated the same scenario 10 times. Simultaneously, we replicated the same scenarios using our simulative approach and show here the closest match (also replicated 10 times) to CDPOP simulations (blue graphs). Five methods for evaluating the genetic makeup are presented. In a), a principal component analysis is applied to the genotype matrix and the differential of the percentage of explained variation by each component is plotted for every replicate. In b), a pairwise Fst analysis between five subpopulations is performed for every replicate and the resulting distribution of Fst is shown. In c), Mantel correlation is calculated between a matrix of genetic and of geographic distances. The resulting Mantel R for every replicate is shown. In d) and e), the allelic frequency of adaptive genotypes is shown as a function of the environmental variables causing selection (representing a case of moderate and strong selection, respectively).

The three ranking coefficients (RMSE, KLD and ΔmR) were scaled using the *scale* R function and averaged, and the resulting value was used to rank simulative variants. In this way, it was possible to find one simulative variant with the best ranking when compared to the CDPOP panmictic population scenario, and another with the best ranking when compared to the CDPOP structured population scenario. These two simulative variants provided the values of *m* and *c* for the simulations on the complete dataset.

Subsequently, we focused on the comparison of the values for the parameters defining the adaptive processes: *s*_*1*_ and *s*_*2*_. For each CDPOP demographic scenario, we searched for the *s*_*1*_ and *s*_*2*_ combination that resulted in a simulative variant that best matched the allelic frequencies of each of the two genotypes implied in selection (moderate and strong). The environmental variable of interest was distributed in 20 equal intervals and within each interval the allelic frequencies of the adaptive genotype were computed. This resulted in the computation of a regression line for each simulation that described the allelic frequency of the adaptive genotype as a function the environmental variable causing the selective constraint (Fig. 2d-e). Next, we calculated the RMSE to compare this regression line between the CDPOP scenarios and the respective simulative variant (*i.e.* those with the optimal *m* and *c* according to the previous analyses) under different *s*_*1*_ and *s*_*2*_ combinations. For the two demographic scenarios, the ranges of *s*_*1*_ and *s*_*2*_ were ranked according to RMSE to represent a moderate to strong selection in the simulations for the complete dataset.

### Sampling design

Four types of sampling design are proposed: three of them differently account for the characteristics of the landscape while one randomly selects the sampling locations. The first is “geographic” (Fig. 3a) and is defined through a hierarchical classification of the sites based on their geographic coordinates. The desired number of sampling locations (L) determines the number of clusters and the geographical center of each cluster is set as a sampling location. The goal of this strategy is to sample sites located as far apart as possible from each other in the geographical space to guarantee spatial representativeness.

**Figure 3.**
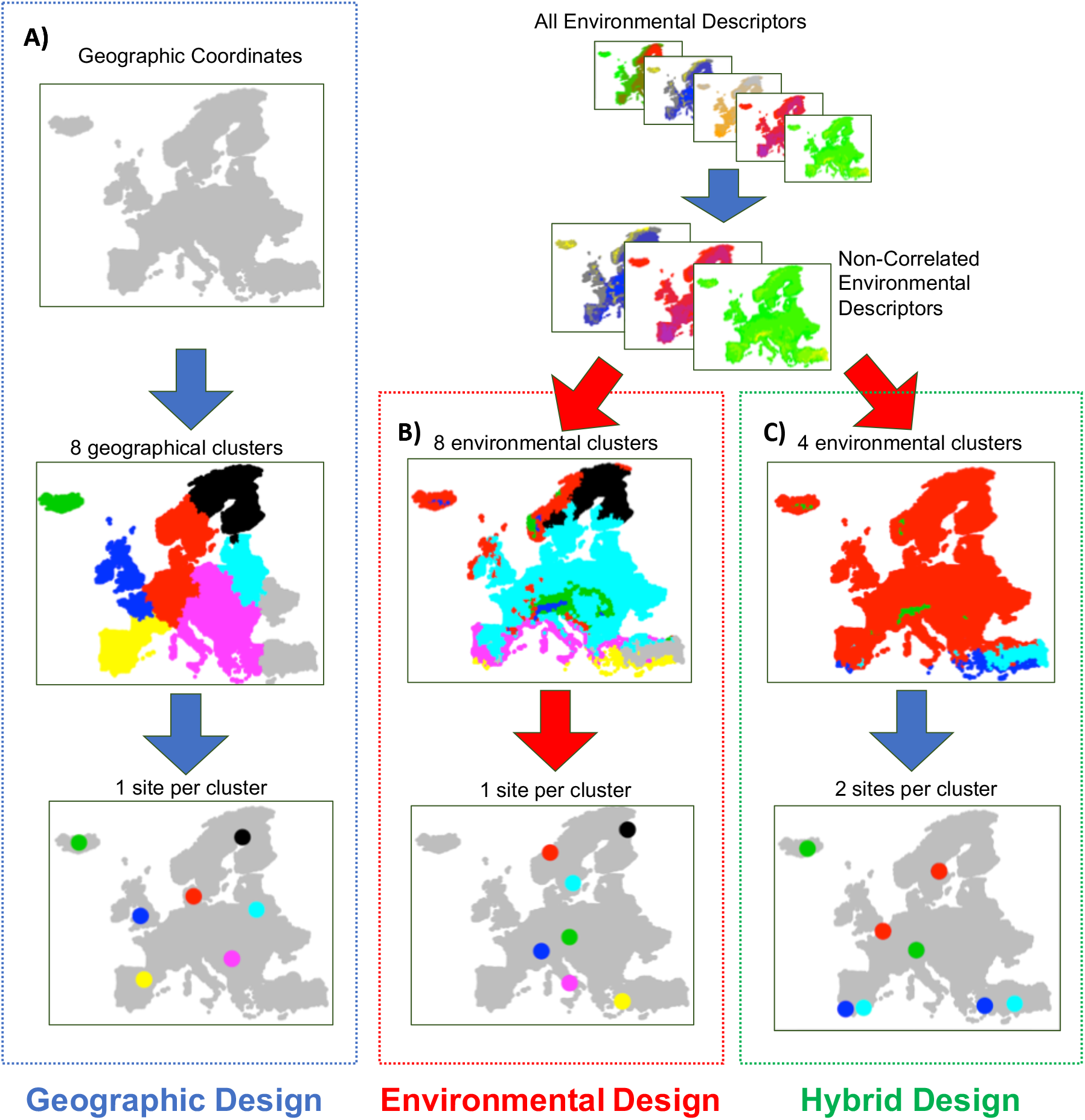
The three sampling design approaches accounting for landscape characteristics. The three maps illustrate how the eight sampling sites are chosen under three different sampling designs. Under a geographic strategy (A), sample location is selected using only geographic coordinates in order to maximize distance between sites. The environmental design (B) is computed using environmental variables (after filtering out highly correlated variables), in order to maximize the climatic distance between the chosen sites. The hybrid strategy (C) is a combination of the first two designs: first the landscape is divided into distinct environmental regions before choosing sites within each region that maximize spatial distance.

The second design type is “environmental” (Fig. 3b). It is based on the computation of distances depending on the values of environmental variables. The latter are first processed by a correlation filter: when two variables are found correlated to each other (R>±0.5), one of them (randomly chosen) is excluded from the dataset. The remaining un-correlated descriptors are scaled (sd=1) and centered (mean=0) using R *scale* function. The scaled values are used to perform a hierarchical clustering between the landscape sites. Like the previous design, the desired number of sampling locations (L) defines the number of clusters. For each cluster, the environmental center is defined by an array containing the mean of the scaled environmental values. Then, the Euclidean distances between this array and the scaled values of each site of the cluster are computed. On this basis, the most similar sites to each center are selected as sampling locations. This strategy aims to maximize environmental contrast between sampling locations and thus favors the detection of adaptive signals (Manel et al., 2012; Riginos et al., 2016).

The third design is “hybrid” (Fig. 3c) and is a combination of the first two. It consists of dividing the landscape into *k* environmental regions and selecting within each of these regions two or more sampling locations based on geographic position. Initially, the environmental variables are processed as for the environmental design (correlation-filter and scaling) and used for the hierarchical classification of the landscape sites. The next step is separating the landscape sites in *k* environmental regions based on this classification. The allowed value of *k* ranges between 2 and half of the desired number of sampling locations (L). We use the R package NbClust (version 3.0, Charrad, Ghazzali, Boiteau, & Niknafs, 2015) to find the optimal value of *k* within this range. The optimal *k* is then used to determine the *k* environmental regions. Next, the number of sampling locations (L) is equally divided across the *k* environmental regions. If *k* is not an exact divisor of L, the remainder of L/*k* is randomly assigned to environmental regions. The number of sampling locations per environment region (L_ki_) can therefore be equal among environmental regions or, at worst, differ by one (for ex. If L=8 and *k*=4: L_k1_=2, L_k2_=2, L_k3_=2, L_k4_=2; if L=10 and *k*=4: L_k1_=3, L_k2_=3, L_k3_=2, L_k4_=2). Sampling locations within environmental regions are chosen based on geographical position. Geographical clusters within each environmental region are formed as in the geographic design, setting L_ki_ as the number of clusters. The landscape site spatially closer to the center of each geographical cluster is selected as sampling location. In such a way, the procedure allows the replication of similar environmental conditions at distant sites, being therefore expected to disentangle neutral and adaptive genetic variation and to promote the detection of variants under selection (Manel et al., 2012; Rellstab et al., 2015; Riginos et al., 2016).

The fourth type of design is “random”: the sampling locations (L) are randomly selected across all the available landscape sites. In our simulations, we tested each type of sampling design with numbers comparable to the ones used in real experiments (see Tab. 1). We used 5 levels of sampling locations L (5, 10, 20, 40 and 50 locations) and 6 of sample sizes N (50, 100, 200, 400, 800 and 1600 individuals). In iterations for which the sample size is not an exact multiple of the number of sites (for ex., 20 sites and 50 individuals), the total number of individuals was changed to the closest multiple (here 40 individuals). The scripts including these procedures were written in R using the functions embedded within the *stats* package (R Core Team, 2016).

### Landscape genomics analysis

We computed association models for each iteration with the SamBada software (version 0.6.0; Stucki et al., 2017). First, the simulated matrix of genotypes is filtered through a customized R function with minor allele frequency <0.05 and major genotype frequency >0.95 to avoid including rare or monomorphic alleles and genotypes, respectively. Secondly, a principal component analysis (PCA) is run on the filtered genotype matrix to obtain synthetic variables accounting for population structure (hereafter referred to as population structure variables; Patterson, Price, & Reich, 2006). The analysis of the eigenvalues of the PCA is carried out in order to assess whether the population structure is negligible for downstream analysis or not (Patterson et al., 2006). At each iteration, the algorithm runs a Tracy-Widom significance test of the eigenvalues, as implemented in the AssocTests R package (version 0.4, Wang, Zhang, Li, & Zhu, 2017). Significant eigenvalues indicate the presence of non-negligible population structure: in these situations, the corresponding principal components will be used as co-variables in the genotype-environment association study.

After filtering, SamBada is used to detect candidate loci for local adaptation. The software is able to run multivariate logistic regression models (Joost et al., 2007) that include population structure as a co-variable, while guaranteeing fast computations (Duruz et al., 2019; Rellstab et al., 2015; Stucki et al., 2017). To ensure compatibility with our pipeline and increase computational speed, we integrated the SamBada method into a customized python script (version 3.5; van Rossum, 1995) based on the Pandas (McKinney, 2010), Statsmodels (Seabold & Perktold, 2010) and Multiprocessing (Mckerns, Strand, Sullivan, Fang, & Aivazis, 2011) packages. *P*-values related to the two statistics (G-score and Wald-score) associated with each association model are computed and subsequently corrected for multiple testing using the R *q-value* package (version 2.6; Storey, 2003). Models are deemed significant when showing a q<0.05 for both tests. When multiple models are found to be significant for the same marker, only the best one is kept (according to the G-score). The pipeline was developed in the R-environment using the *stats* library.

### Simulations and evaluation of the performance

Each combination of demographic scenarios, sampling designs, number of sampling locations and sample sizes was replicated 20 times for a total of 4,800 iteration (Tab. 2). A new genetic matrix was randomly redrawn for each iteration to change the selective forces implying local adaptation and the demographic set-up determining the neutral loci. At the end of each iteration, three diagnostic parameters were computed:

- True Positive Rate of the analysis (TPR or statistical power): percentage of true associations detected to be significant;
- False Discovery Rate of the analysis (FDR): percentage of false association among the significant ones;
- Positive Predictive Value (PPV; Marshall, 1989) of the ten strongest associations: significant associations were sorted according to the association strength (*β*, the value of the parameter associated to environmental variable in the logistic model). PPV represents the percentage of true associations among the best ten associations according to *β*.

**Table 2.** Table of factors varying in the simulative approach. Two different demographic scenarios are possible, one in which there is no neutral genetic structure (panmictic population) and one in which there is a structured variation (structured population). We then used sampling strategies emulating those observed in real experiments. Three different sampling design approaches accounting for landscape characteristics are proposed: one maximizing the spatial representativeness of samples (geographic), one maximizing the environmental representativeness (environmental) and one that is a combination of both (hybrid). A fourth sampling design picks sampling locations randomly. The numerical ranges we employed were comparable to those from real experiment: 5 levels for number of sampling locations spanning from 5 to 50 sites, and 6 levels of sample sizes (i.e. total number of samples) from 50 to 1600 samples. For each combination of the aforementioned factors, 20 replicates were computed differing in the number and types of selective forces driving adaptation. In total, 4,800 simulation were computed.

| Factor | # levels | Levels |
| --- | --- | --- |
| Demographic Scenarios | 2 | Panmictic Population, Structured Population |
| Sampling Design (*D*) | 4 | Geographic, Environmental, Hybrid, Random |
| Sampling Locations (*L*) | 5 | 5, 10, 20, 40, 50 |
| Sampling Size (*N*) | 6 | 50, 100, 200, 400, 800, 1600 |
| Replicates | 20 |  |
| Total | 4800 |  |

After the simulations, an analysis of ranks (Kruskal-Wallis test; Kruskal & Wallis, 1952) was performed using the *kruskal.test* R function to test whether TPR, FDR and PPV were significantly influenced (p<0.01) by each of the three elements underlying the sample strategy (*i.e.* sampling design, number of sampling locations and sample size; Tab. 2). Contextually, we computed the epsilon-squared (E^2^) coefficient (as implemented in the *rcompanion* R package, version 2.2.1; Mangiafico, 2019) that quantifies, on a scale from 0 to 1, the influence of each sampling element on the three diagnostic parameters (Tomczak & Tomczak, 2014). Finally, we calculated the changes in the median values of TPR, FDR and PPV across the levels of each element underlying the sample strategy. In the case of numerical elements (*i.e.* number of sampling locations and sample size), we quantified the changes in TPR, FDR and PPV along with the increments of the ordinal factor levels (for ex.: the TPR median increase between sample sizes of 100 to 200, 200 to 400, 400 to 800, etc.). In the case of sample design, where the factor levels are not ordinal, we compared each design approach against a random sampling scheme.

## Results

### Parameters of simulations

For the panmictic population scenario, the simulative variant best matching the CDPOP results was obtained with the coefficients *m* = 1 and *c* = 0.5, whereas for the structured population scenario, the simulative variant was best at *m* = 10 *and c* = *Unif*(0.2, 0.4) (Fig. 2a-c, Box S2, Tab. S2a-b). In the panmictic population scenario, we found that the moderate selection case was best emulated by *s*_*1*_ = 0.4 and *s*_*2*_ = −0.2 and the strong selection by *s*_*1*_ = 0.3 and *s*_*2*_ = +0.1. In the structured population scenario, the moderate selection found its best match in the simulative variant with *s*_*1*_ = 0 and *s*_*2*_ = −0.1 while the strong selection in the one set with *s*_*1*_ = 0 and *s*_*2*_ = +0.2 (Fig. 2d-e, Box S2, Tab. S2c-d).

### True Positive Rate

In general, the panmictic population scenario simulations showed higher TPR (Mdn_PAN_=40% [IQR=0-90%]) than simulations performed under the structured population scenario (Mdn_STR_=0% [IQR=0-40%]; Fig. 4a-c). For both scenarios, the main influence on TPR was found to be sample size (E^2^_PAN_=0.815, E^2^_STR_=0.613; Tab. 3c). Smaller sample sizes (N= 50, 100) resulted in TPR close or equal to zero for both demographic scenarios (Fig. 4c, Tab. S3c). Under the structured population scenario, an increase of TPR started from N=200 (Tab. S3c), leading to an initial increase of 5% of the median TPR for every 10 additional samples. At N=400, this increment progressively became less abrupt until reaching a maximal value at N=800 (Mdn=100% [IQR=60-100%]; Fig. 4c; Tab. S3c). By comparison, the panmictic population scenario showed an increase in TPR starting at N=400, with a more constant and less abrupt rate of increase (Fig. 4c, Tab. S3c). Under this scenario, a N=1600 was not sufficient to yield maximal TPR (Mdn=80% [IQR=60-90%]; Fig. 4c).

**Table 3.** Results of Kruskal-Wallis (KW) rank analysis. The table shows the epsilon-squared (E^2^) coefficient associated to the KW test for the three diagnostic parameters of the analysis (TPR: true positive rate, a-c; FDR: false discovery rate, d-f; PPV: positive predictive value of among the ten strongest significant association models, g-i) for every element determining the sampling strategy (sampling design: a, d, g; number of locations: b, e, h; sample size: c, f, i) under the two demographic scenarios, panmictic and structured population. E^2^ ranges between 0 and 1, where the higher the value the stronger the sampling strategy element drives the differences in the diagnostic parameter. The asterisks represent the respective degree of significance of the KW test (*: p<0.01, **: p<0.01, ***: p<0.001, **** p<0.0001, ***** p<0.00001).

| TPR |  |  |  |  |  |
| --- | --- | --- | --- | --- | --- |
| a) Sampling Design |  | b) # of Sampling Locations |  | c) Sample Size |  |
| Panmictic | Structured | Panmictic | Structured | Panmictic | Structured |
| 0.0163***** | 0.0229***** | 0.00766* | 0.17***** | 0.815***** | 0.613***** |
| FDR |  |  |  |  |  |
| d) Sampling Design |  | e) # of Sampling Locations |  | f) Sample Size |  |
| Panmictic | Structured | Panmictic | Structured | Panmictic | Structured |
| 0.00122 | 0.00733** | 0.0116*** | 0.127***** | 0.621***** | 0.408***** |
| PPV |  |  |  |  |  |
| g) Sampling Design |  | h) # of Sampling Locations |  | i) Sample Size |  |
| Panmictic | Structured | Panmictic | Structured | Panmictic | Structured |
| 0.00226 | 0.0264***** | 0.0124**** | 0.19***** | 0.63***** | 0.381***** |

**Figure 4.**
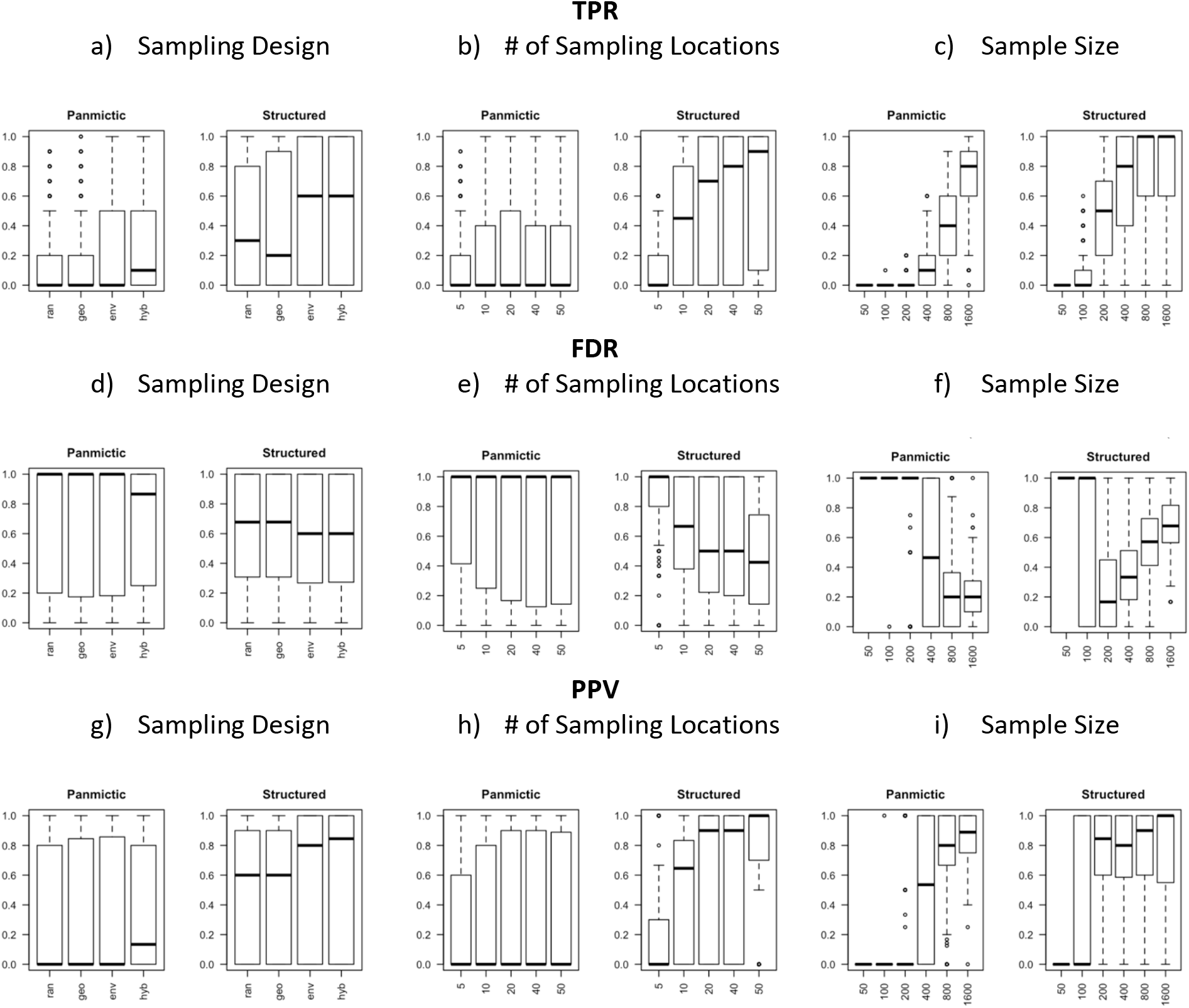
Effects of sampling strategy on the landscape genomics simulations. The plots display how the performance of landscape genomics experiments is driven by changes in the elements defining the sampling strategy. Three diagnostic parameters are used to measure the performance of each strategy: true positive rate (TPR; a-c) and false discovery rate (FDR; d-for the analysis and the positive predictive value of the ten strongest significant association models (PPV; g-i). For each diagnostic parameter, we show the effect of sampling design (a, d, g; ran=random, geo=geographic, env=environmental, hyb= hybrid), number of sampling locations (b, e, h; 5, 10, 20, 40 or 50 sites) and sample size (c, f, I; 50, 100, 200, 400, 800, 1600 individuals) under two demographic scenarios: panmictic and structured population.

The effect of number of sampling locations on TPR was significant, even though weaker than the effect of sample size (E^2^_PAN_=0.008, E^2^_STR_=0.17; Tab. 3a; Fig. 4b). Under the panmictic population scenario in particular, an increase in the number of sampling locations did not change the median TPR, but its inter-quartile range (Fig. 4b, Tab. S3b). Conversely, TPR was affected by the number of sampling locations under the structured population scenario (Fig. 4b, Tab. S3b). This effect was particularly evident between L=5 and L=10, where additional sampling sites led to an increase of the median TPR by 10% (Tab. S3b). At higher numbers of sampling sites (L=20, 40 and 50) the incremental rate of TPR was less evident but still positive (Tab. S3b).

Similar to the influence of sampling locations, the type of sampling design had a minor effect on TPR when compared to the effect that sample size had (E^2^_PAN_=0.0163, E^2^_STR_=0.1229; Tab. 3a; Fig. 4a). When compared to the random approach, a hybrid design approach was seen to increase the median TPR by 10% and 30% under panmictic and structured population scenarios, respectively (Fig. 4a, Tab S3a). The two other design approaches only affected median TPR for the structured population scenario; compared to a random sampling scheme, the environmental design increased TPR by 30%, while the geographical design decreased TPR by 10% (Fig. 4a, Tab. S3a).

### False Discovery Rate

False discoveries generally appeared at a higher rate under a panmictic population scenario (Mdn_PAN_=100% [IQR=20-100%]) than under a structured population scenario (Mdn_STR_=63% [IQR=20-100%]; Fig. 4d-f). Sample size had the greatest effect on FDR for both population scenarios (E^2^_PAN_=0.621, E^2^_STR_=0.408; Tab. 3f; Fig. 4f). For the panmictic population scenario, median FDR was 100% at smaller sample sizes (N=50, 100 and 200; Fig. 4f), but between N=200 and N=400, the FDR began to decrease by 2.6% for every ten additional samples taken (Tab. S3c). The reduction in FDR was less abrupt after N=400, and null after N=800 (Tab. S3c). At N=1600, median FDR was 20% [IQR=10-30%] (Fig. 4f). The structured population scenario produced a different pattern: the largest median FDR was found at smaller sample sizes (N=50 and 100), before a steep decrease was observed closer to N=200 (Fig. 4f, Tab. S3c). At larger sample sizes (N=400, 800, 1600), FDR showed a logarithmic increase in growth rate where, at its most abrupt (between N=200 and 400), there was an increase of 0.8% FDR for every ten additional samples (Fig. 4f, Tab. S3c). For the structured population scenario, N=1600 resulted in a median FDR of 68% [IQR=57-82%].

The effect of sampling location number on FDR was significant, albeit weaker than the effect of sample size on FDR, under both population scenarios (E^2^_PAN_=0.012, E^2^_STR_=0.127; Tab. 3e, Fig. 4e). Similar to the pattern for TPR, the number of locations sampled under the panmictic population scenario did not alter the median FDR, but rather its inter-quartile range (Fig. 4e). By contrast, the structured population scenario showed a decrease of median FDR along with the increment of the number of sampling locations (Fig. 4e, Tab. S3b). This decrease was more abrupt increasing from 5 to 10 sampling locations, where additional sites led to a median FDR reduction of 6%, than between higher numbers of sampling locations (L=20, 40, 50; Tab. S3b).

Sampling design only showed a significant effect on false discovery rates under the structured population scenario, but it was not as strong as the influences of sample size and sampling locations (E^2^_STR_=0.007; Tab. 3d, Fig. 4d). When compared to a random sampling scheme, both the environmental and hybrid sampling schemes showed a decrease in median FDR of 7%, while the geographic scheme showed a slight increase in FDR (+0.05%; Fig. 4d, Tab. S3a).

### Positive Predictive Value

The PPVs of the ten strongest significant associations (hereinafter simply referred to as PPV) was generally higher under the structured population scenario (Mdn_PAN_=70% [IQR=0-100%]) than under the panmictic population scenario (Mdn_PAN_=0% [IQR=0-80%]; Fig. 4g-i). As with TPR and FDR, sample size had the strongest influence on PPV under both population scenarios (E^2^_PAN_=0.63, E^2^_STR_=0.381; Tab. 3i, Fig. 4i). Under the panmictic population scenario, median PPV was 0% for the smaller sample sizes (N=50, 100 and 200; Fig. 4i), after which patterns of increase were observed: from N=200 to 400 there was an increase of PPV of 2.6% for every 10 additional samples, and from N=800 to 1600 PPV continued to increase though it was less abrupt, resulting in a median PPV of 88% [IQR=75-100%] at N=1600 (Fig. 4i, Tab. S3c). Under the structured population scenario, fewer samples were required to observe a similar increment: while median PPV was 0 for N=50 and N=100, from N=100 to N=200 the median PPV increased by 8.4% for every ten additional samples (Fig. 4i, Tab. S3c). The increment of PPV became gradually weaker when transitioning between higher levels (N=400, 800 and 1600) and led to a median PPV of 100% [IQR=57.5-100%] at N=1600.

Similar to TPR and FDR, the effect of sampling location number on PPV was significant but weaker than when compared to the effect of sample size (E^2^_PAN_=0.0124, E^2^_STR_=0.19; Tab. 3h, Fig. 4h). This effect was particularly evident under the structured population scenario, where an increase of the number of sampling locations strongly raised median PPV (Fig. 4h). The strongest PPV increment was observed between L=5 and 10, where each additional sampling location raised the median PPV by 13% (Fig. 4h, Tab. S3b). With more sampling locations (L=20, 40 and 50) the rate of increase of PPV remained but was weaker (Fig. 4h, Tab. S3b). In the panmictic population scenario, an increase in the number of sampling locations affected the inter-quartile range of PPV in particular, but not the medians (Fig. 4h).

The sampling design used resulted in rate changes for PPV under the structured population scenario, despite being less strong than when compared to the other elements (E^2^_STR_=0.0264; Tab. 3g, Fig. 4g). When compared to a random sampling scheme, the hybrid design and the environmental design increased the median PPV by 24% and 20% respectively, while the geographic design did not result in any changes (Fig. 4g, Tab. S3a).

## Discussion

The simulations presented in this study highlight that sampling strategy clearly drives the outcome of a landscape genomics experiment, and that the demographic characteristics of the studied species can significantly affect the analysis. Despite some limitations that will be discussed below, the results obtained make it possible to answer three questions that researchers are confronted with when planning this type of research investigation.

### How many samples are required to detect any adaptive signal?

In line with the findings of previous studies (e.g. Lotterhos & Whitlock, 2015), our results suggest that sample size is the key factor in securing the best possible outcome for a landscape genomics analysis. Where statistical power is concerned, there is an unquestionable advantage in increasing the number of samples under the scenarios tested. When focusing on the panmictic population scenario, we found a lack of statistical power in simulations for N≤200, while detection of true positives increased significantly for N≥400 (Fig. 4c). As we progressively doubled sample size (N=800, 1600), TPR linearly doubled as well (Fig. 4c). Under the structured population scenario, this increase in statistical power started at N≥100 and followed a logarithmic trend that achieved the maximum power at N≥800 (Fig. 4c).

These results show that it is crucial to consider the population’s demographic background to ensure sufficient statistical power in the analyses, as advised by several reviews in the field (Balkenhol et al., 2017; Manel et al., 2012; Rellstab et al., 2015). In fact, the allelic frequencies of adaptive genotypes respond differently to a same environmental constraint under distinct dispersal modes (Fig. 2d-e). When individual dispersal is limited by distance (structured population scenario), the allelic frequencies of adaptive genotypes are the result of several generations of selection, resulting in a progressive disappearance of non-adaptive alleles from areas where selection acts. When the dispersal of individuals is completely random (panmictic population scenario), the same selective force only operates within the last generation, such that even non-adaptive alleles can be found where the environmental constraint acts. Under these premises, a correlative approach for studying adaptation (such as SamBada) is more likely to find true positives under a structured population scenario rather than under a panmictic one.

The dichotomy between structured and panmictic populations also emerges when analyzing false discovery rates. Under the panmictic population scenario, increasing the number of individuals sampled reduced FDR, while the inverse pattern was seen under a structured population scenario (Fig. 4f). The issue of high false positives rates under structured demographic scenarios is well acknowledged in landscape genomics (De Mita et al., 2013; Rellstab et al., 2015). Population structure results in gradients of allele frequencies that can mimic and be confounded with patterns resulting from selection (Rellstab et al., 2015). As sample size increases, the augmented detection of true positives is accompanied by the (mis-)detection of false positives. Under the panmictic population scenarios, these confounding gradients of population structure are absent (Fig. 2a-c) and high sample sizes accentuate the detection of true positives only (Fig. 4f).

Working with FDR up to 70% (Fig. 4f) might appear excessive, but this should be contextualized in the case of landscape genomics experiments. The latter constitute the first step toward the identification of adaptive loci, which is generally followed by further experimental validations (Pardo-Diaz, Salazar, & Jiggins, 2015). Most landscape genomics methods test single-locus effects (Rellstab et al., 2015). This framework is efficient for detecting the few individual loci that provide a strong selective advantage, rather than the many loci with a weak individual-effect (for instance those composing a polygenic adaptive trait; Pardo-Diaz et al., 2015). For this reason, when researchers are faced with a high number of significant associations, they tend to focus on the strongest ones (Rellstab et al., 2015), as we did here by measuring the PPV of the ten strongest associations. By relying on this diagnostic parameter, we could show that increasing sample size ensures that the genotypes more strongly associated with environmental gradients are truly due to adaptive associations (Fig. 4i). Under these considerations, acceptable results are obtainable with moderate sample sizes: a median PPV of at least 50% was found with simulations with N=400 and N=200 under panmictic and structured population scenario, respectively.

Each landscape genomic experiment is unique in terms of environmental and demographic scenarios, which is why it is not possible to propose a comprehensive mathematical formula to predict the expected TPR, FDR and PPV based solely on sample size. When working with a species with a presumed structured population (for instance, wild land animals), we advise against conducting experiments with fewer than 200 sampled individuals, as the statistical requirements to detect true signals are unlikely to be met. Panmixia is extremely rare in nature (Beveridge & Simmons, 2006), but long-range dispersal can be observed in many species such as plants (Nathan, 2006) and marine organisms (Riginos et al., 2016). When studying species of this kind, it is recommendable to increase sample size to at least 400 units.

### How many sampling sites?

Increasing the number of samples inevitably raises the cost of an experiment, largely resulting from sequencing and genotyping costs (Manel et al., 2010; Rellstab et al., 2015). Additionally, field work rapidly increases the cost of a study in cases where sampling has to be carried out across landscapes with logistic difficulties and physical obstacles. Therefore, it is both convenient and economical to optimize the number of sampling locations to control for ancillary costs.

De Mita et al. (2013) suggested that increasing the number of sampling locations would raise power and reduce false discoveries. The present study partially supports this view. A small number of sampling locations (L=5) was found to reduce TPR and PPV while increasing FDR, compared to using more sampling locations (L=10, 20, 40 and 50; Fig. 4b, e, h). This is not surprising, because when sampling at a small number of locations the environmental characterization is likely to neglect some contrasts and ignore confounding effects between collinear variables (Leempoel et al., 2017; Manel et al., 2010). This was particularly evident under the structured population scenario (Fig. 4b, e, h). In contrast, we found that higher numbers of sampling locations (L=40 and 50) provided little benefits in terms of TPR, FDR and PPV, compared to a moderate number of locations (L= 20; Fig. 4b, e, h). These discrepancies with previous studies are probably due to differences in the respective simulative approaches applied (we use several environmental descriptors instead of one) and the characteristics of the statistical method we employed to detect signatures of selection. In fact, as a number of sampling locations is sufficient to portray the environmental contrasts of the study area, adding more locations does not bring additional information and therefore does not increase statistical power. The implications of the information described above are considerable since the cost of field work can be drastically reduced with marginal countereffects on statistical power and false discoveries.

### Where to sample?

Compared with random or opportunistic approaches, sampling designs based on the characteristics of the study area are expected to improve the power of landscape genomics analysis (Lotterhos & Whitlock, 2015). We developed three distinct methods to choose sampling locations accounting for geographical and/or environmental information (geographic, environmental and hybrid designs). We confronted these design approaches between themselves and with random sampling schemes. The approach based on geographic position (geographic design) resulted in statistical power similar to the random designs (Fig. 4a, d, f), while those based on climatic data (environmental and hybrid design) displayed remarkably higher TPRs and PPV and slightly lower FDR (Fig. 4a, d, f). These beneficial effects on the analysis were accentuated under the structured demographic scenario.

These results match previous observations: methods conceived to take advantage of environmental contrasts facilitate the detection of adaptive signals (Manel et al., 2012; Riginos et al., 2016). Furthermore, the hybrid design prevents the sampling of neighboring sites with similar conditions, therefore avoiding the superposition between adaptive and neutral genetic variation (Manel et al., 2012). This is likely to explain why the hybrid design slightly outperformed the environmental approach (Fig. 4a, d, f). For these reasons, we strongly advise in using a sampling scheme accounting for both environmental and geographical representativeness. Without bringing any additional cost to the analysis, this approach can boost statistical power of up to 30% under a complex demographic scenario (Tab. S3a), in comparison to a regular (geographic) or random sampling scheme.

### Limitation

The preliminary run of comparison with a well-established forward-in-time simulation software (CDPOP) displayed the pertinence of our customized simulative approach (Fig. 2). The neutral genetic variation appeared as random under the panmictic population scenario (no skew on the PC graph, Fst close to 0, mR close to 0) and structured under the structured population scenario (skew in the PCA graph, Fst higher than 0, mR different from 0; Fig. 2a-c). Adaptive allele frequencies also matched theoretical expectations: the responses along the environmental gradients were more stressed under the structured population scenario than under the panmictic one (Fig. 2d-e).

Nonetheless, the use of *forward-in-time* simulations on the complete dataset (used by De Mita et al., 2013; Lotterhos & Whitlock, 2015) would probably have resulted in more realistic scenarios. In order to be used in a framework as the one proposed here, the *forward-in-time* methods should be compatible with a large number of spatial locations (*i.e.* potential sampling sites), hundreds of individuals per location and a genetic dataset counting at least one thousand loci, of which 10 set as adaptive against distinct environmental variables. Importantly, all these requirements should be fulfilled at a reasonable computational speed (with our method, for instance, genotypes are computed in a few seconds). As far as we know, there are no existing software meeting these criteria.

## Conclusions

The present work provides guidelines for optimizing the sampling strategy in the context of landscape genomic experiments. Our simulations highlight the importance of considering the demographic characteristic of the studied species when deciding the sampling strategy to be used. For species with limited dispersal, we suggest working with a minimum sample size of 200 individuals to achieve sufficient power for landscape genomic analyses. When species display long-range dispersal, this number should be raised to at least 400 individuals. The costs induced by a large number of samples can be balanced by reducing those related to field work. In cases where a moderate number of sampling locations (20 sites) is sufficient to portray the environmental contrasts of the study area, there is only minimal statistical benefit for sampling a larger number of sites (40 or 50). Furthermore, we describe an approach for selecting sampling locations while accounting for environmental characteristics and spatial representativeness, and show its benefic effects on the detection of true positives.

## Acknowledgements

We thank the anonymous reviewers for the useful comments and suggestions provided during the redaction of this paper. We acknowledge funding from the IMAGE (Innovative Management of Animal Genetic Resources) project funded under the European Union’s Horizon 2020 research and innovation program under grant agreement No. 677353.

## Data Accessibility Statement

All scripts and data used to perform this analysis are publicly available on Dryad (doi:10.5061/dryad.m16d23c).

## Author Contributions

OS and SJ designed research, OS performed research, OS, EV, AG, ER and SJ analyzed the results and wrote the paper. All the authors undertook revisions, contributed intellectually to the development of this manuscript and approved the final manuscript.

## Supplemental Information

**Supplementary Tab. 1.**
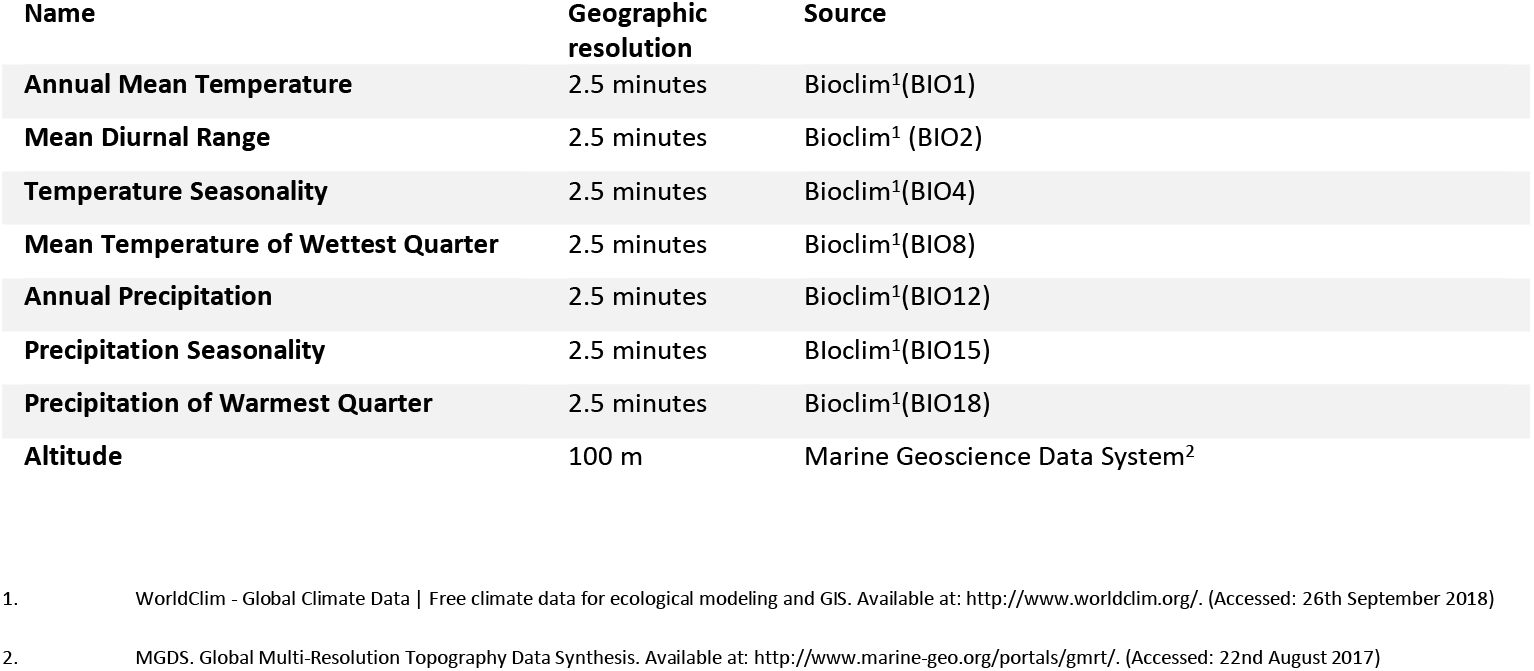
List of environmental variables employed.

**Supplementary Box 1.**
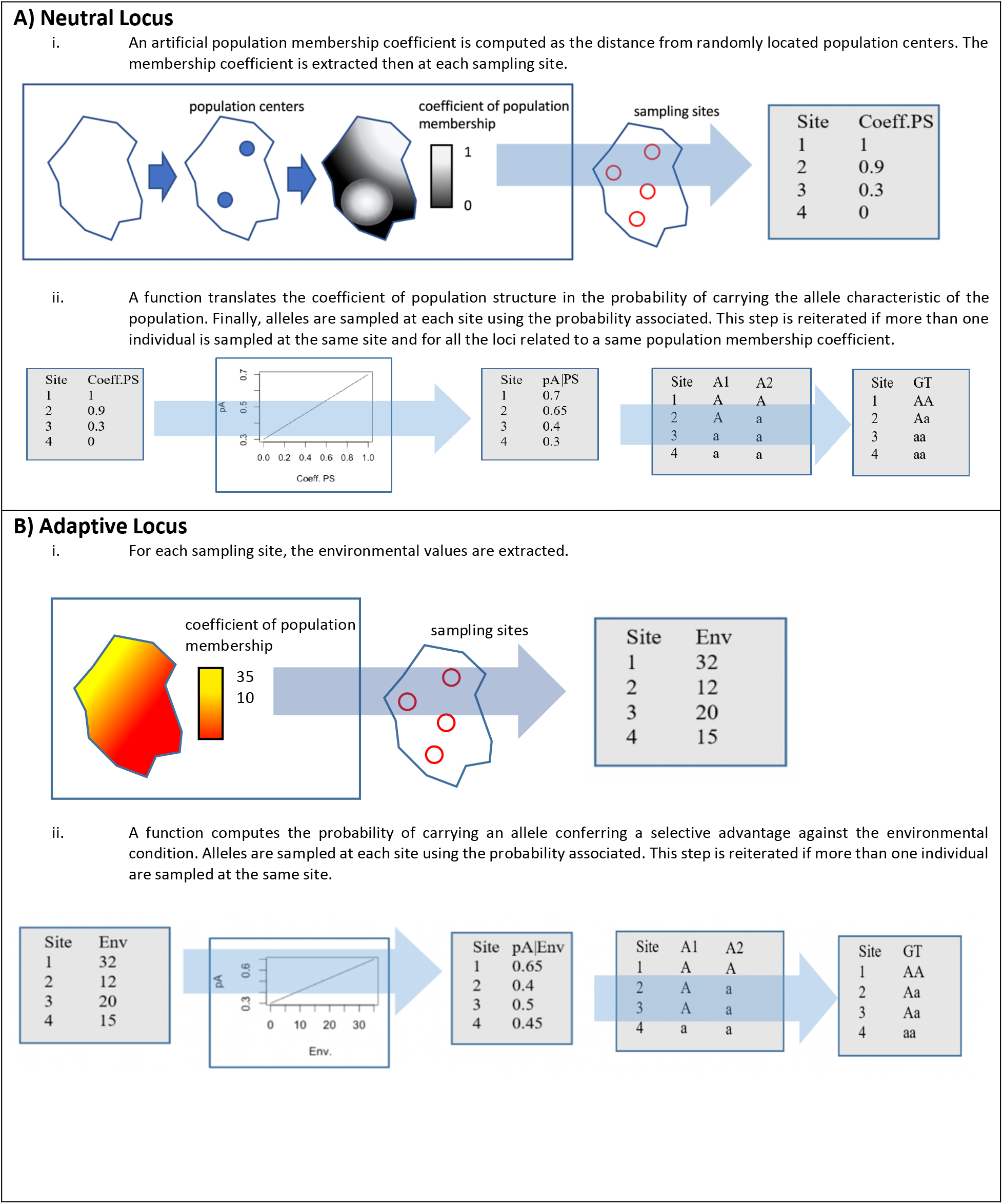
Computation of the genotype matrix. The vignettes describe how genotypes were computed during simulations. At each iteration, a new genotype matrix counting 1’000 loci was generated. Ten of them were set as adaptive and followed the respective pipeline, while the others were set as neutral and computed accordingly.

---

**Supplementary Box 2. Formulae and parameters for genotype computations**

The probability function for the allele A depending on a population membership coefficient is calculated as follows:

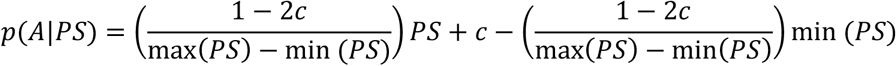

where *PS* is a population membership coefficient and *c* a parameter representing the strength of the relationship. This parameter can range between 0 (strongest relation, *i.e.* maximal and minimal *PS* returns *p*=1 and *p*=0, respectively) and 0.5 (no relation, any level of *PS* returns *p*=0.5).

Similarly, probability for the allele A depending on environmental selection is calculated as follows:

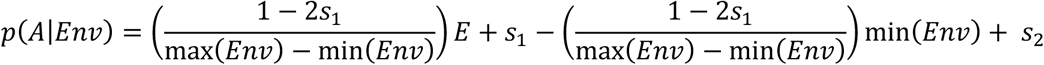

where *Env* are the values of the environmental variable and *s_1_* represents the strength of selection and behaves as the *c* in the previous equation. The additional parameter *s*_*2*_ provides a baseline of allele frequency.

In our simulations, we set two scenarios employing the following parameters:

- *panmictic population scenario* (random neutral structure): *c=0.5, s_1_=Unif(0.3, 0.4), s_2_=Unif(−0.2,0.1)*
- *structured population scenario* (strong population structure): *c=Unif(0.2,0.4), s_1_=0, s_2_=Unif(−0.1,0.2)*

**Supplementary Figure 1.**
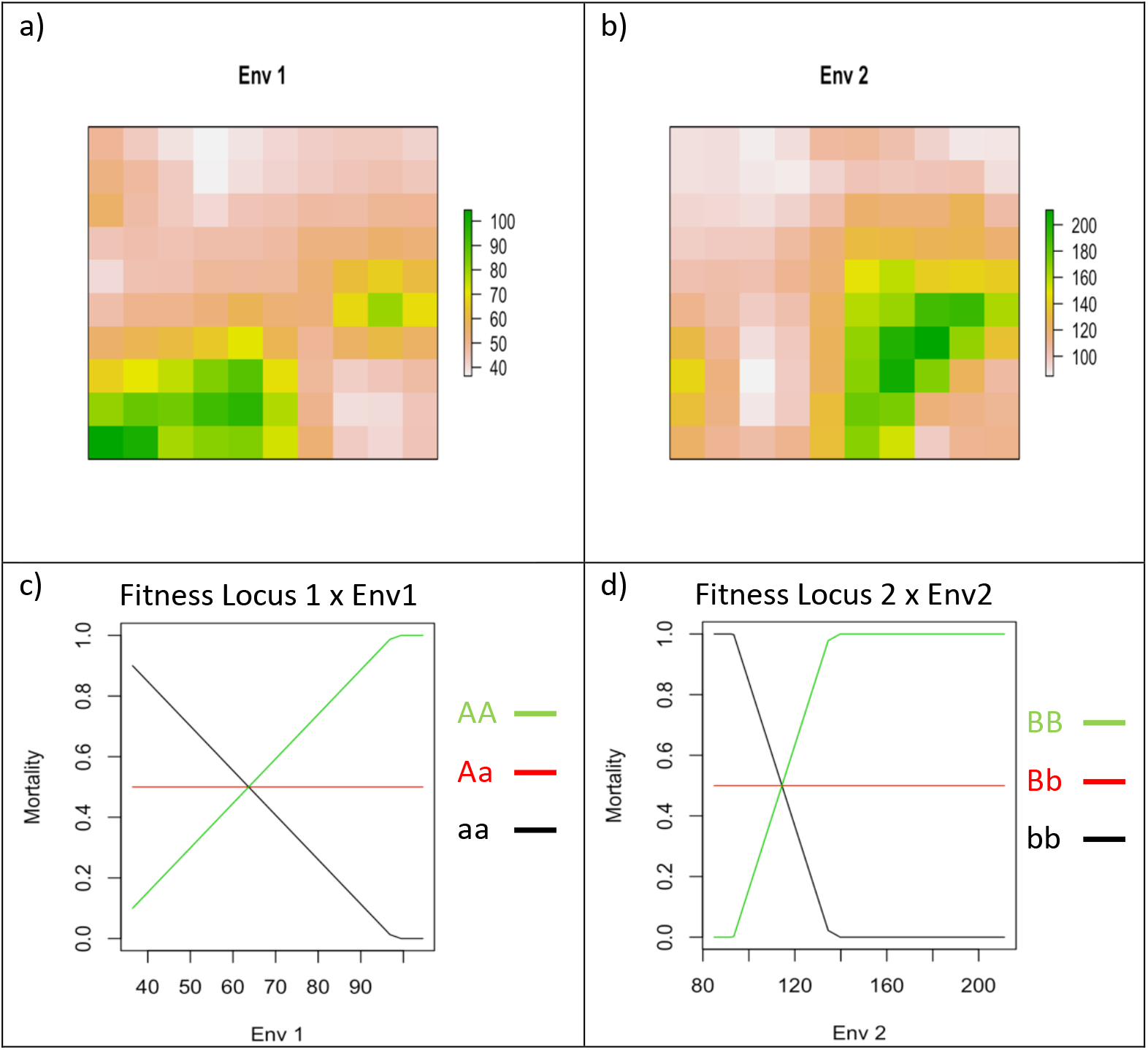
Environmental gradients and fitness constraint employed in the CDPOP validation run. Panel a) and b) show the distribution of the two environmental variables across the 10-by-10 cells grid used for the CDPOP simulation. Plots in panels c) and d) show the fitness constraint set for the two environmental variables and how the respective adaptive genotypes modulate mortality.

**Supplementary Table S2.**
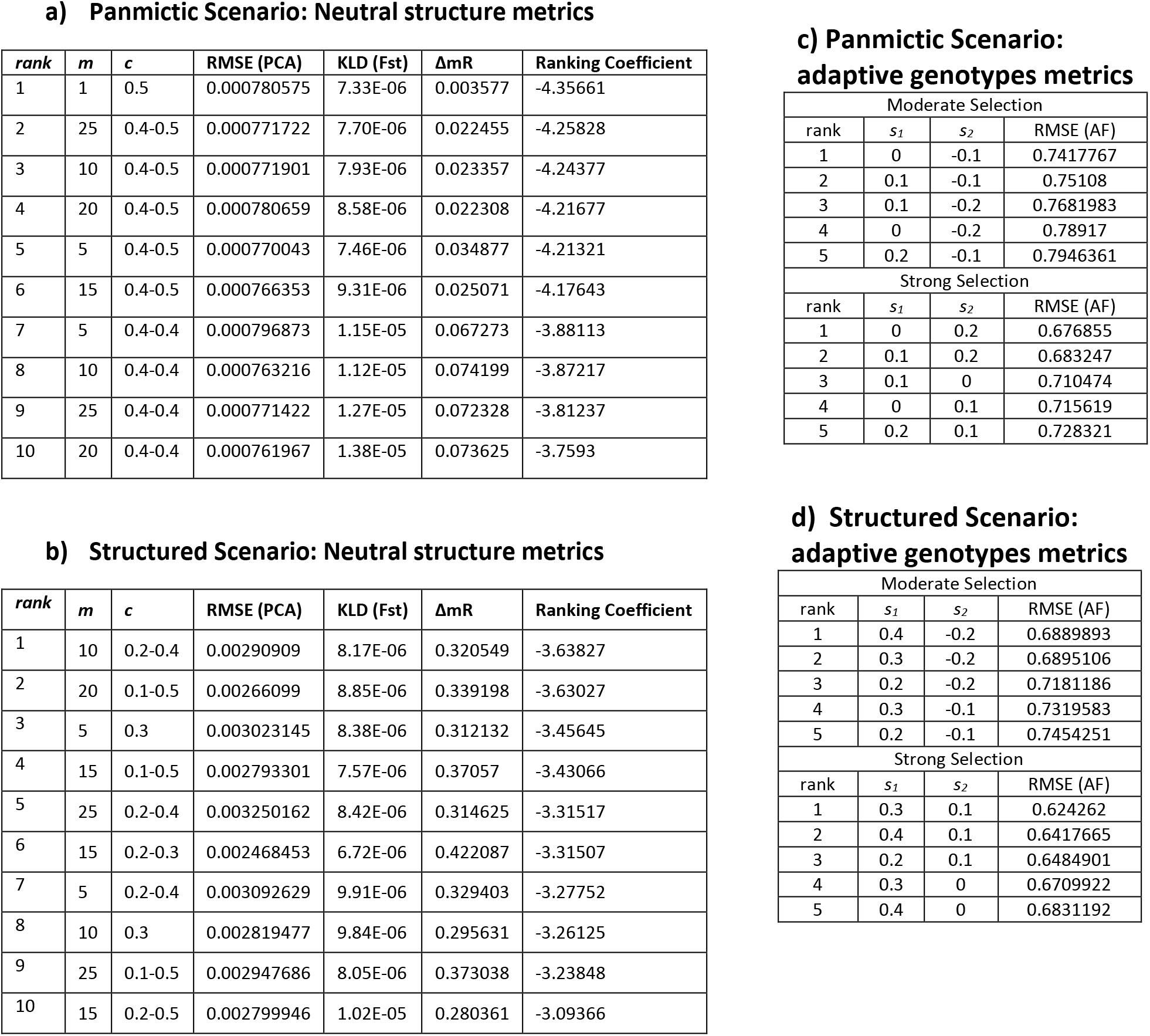
CDPOP vs. our simulative approach comparison metrics. The tables show the rank of the simulative variants computed with our method (and defined by parameters *m, c s*_*1*_ and *s*_*2*_) that best matched the CDPOP replicates. In a) and b) are shown the metrics used to compare the neutral genetic structure with the CDPOP case of a panmictic population and a structured population, respectively. The three metrics employed are 1) the average random mean squared error (RMSE) when comparing the curves describing the differential of explained variation by the genetic principal components; 2) the Kullback-Leibler Divergence (KLD) used to compare the pairwise-Fst distributions; 3) the difference in the average mantel correlation (ΔmR), which describes the link between genetic and geographic distances. The ranking coefficient is the sum of the three scaled metrics. In c) and d) the comparison concerns the adaptive genotypes computed in panmictic structured scenario of CDPOP, respectively. Here the RMSE compares, for our simulation and CDPOP runs, the allelic frequency of the adaptive genotype as a function of the environmental variable causing adaptation

**Supplementary Table S3.**
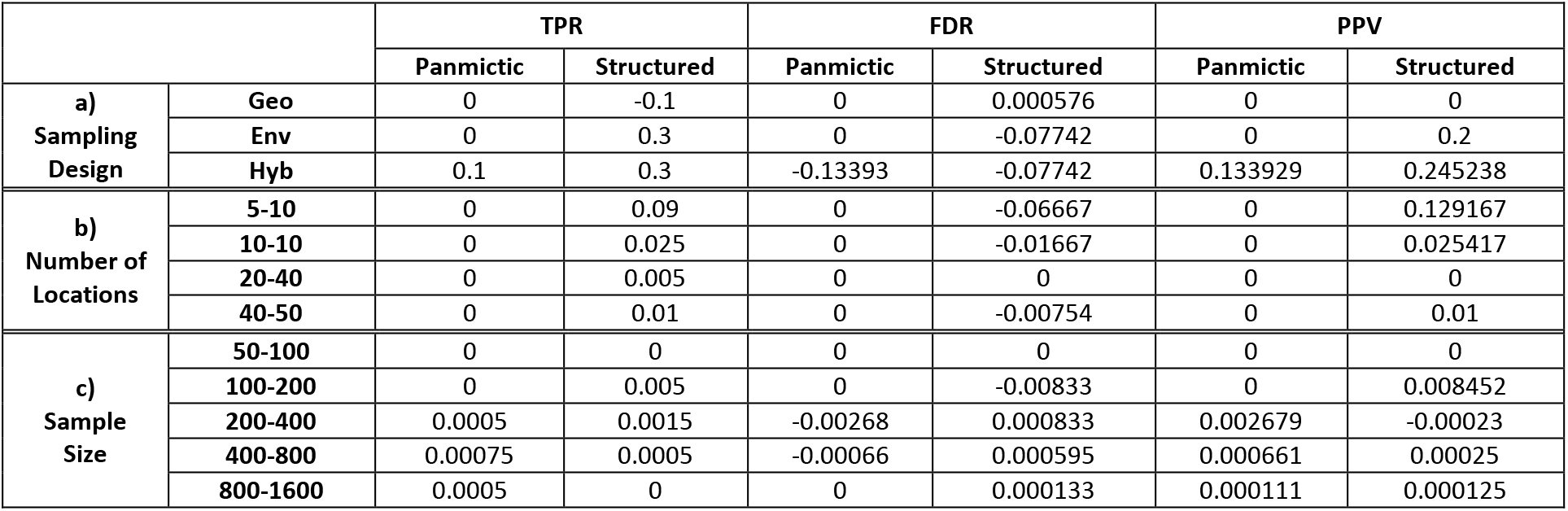
Changes in the analysis results under different sampling strategy. The table shows the changes in the median value of the three diagnostic parameters (TPR, FDR and PPV) used to evaluate the performance of the landscape genomics analysis. In a) the changes concern the different sampling design approaches (geo: geographic, env: environmental, hyb: hybrid) as compared to the random one. In b), the comparison focuses on the number of sampling locations showing, for a given range of locations, by how much an additional sampling site increases the median of the diagnostic parameter. In c) is shown, for a given interval of sample size, by how much the median of the diagnostic parameter is increased by an additional sample. The results for the two demographic scenarios, panmictic and structured, are shown separately.

